# Visualizing early *Mycobacterium tuberculosis* interactions with murine lung macrophages using intravital imaging

**DOI:** 10.64898/2026.03.04.708340

**Authors:** Yookyung Jung, Bing Chen, Catherine Vilchèze, William R. Jacobs, David Entenberg

## Abstract

Intravital microscopy enables direct visualization of dynamic cellular processes within intact tissues, but its application to *Mycobacterium tuberculosis* (Mtb) has been limited by Biosafety Level 3 (BSL-3) containment requirements and the technical challenges of stabilizing the lung for high-resolution imaging. Here, we present a protocol that combines the thoracic Window for High-Resolution Imaging of the murine Lung (WHRIL) with a genetically defined, triple-auxotrophic Mtb strain (mc^2^7902) approved for use under BSL-2 conditions. We describe the construction of a tdTomato-expressing derivative (mc^2^8471) preparation of bacteria for intravenous infection and intravital imaging in reporter mice. This system enables visualization of rapid bacterial entry into the pulmonary vasculature, subsequent aggregation, and vascular occlusion, dissemination into the lung parenchyma, and macrophage uptake over three days post-infection. This protocol provides the first practical platform for real-time intravital imaging of mycobacteria in the lung and establishes a foundation for mechanistic studies of bacterial physiology, host recognition, and immune-mediated clearance using safe Mtb surrogates.

**Summary:** This protocol describes a biosafety level 2 (BSL-2)-compatible intravital imaging platform for visualizing *Mycobacterium tuberculosis* (Mtb) in the intact murine lung at single cell resolution. By combining the Window for High-Resolution Imaging of the murine Lung (WHRIL) with a fluorescently labeled, genetically defined triple auxotrophic Mtb strain (mc^2^7902), this approach overcomes long-standing biosafety and technical barriers that have prevented real-time imaging of mycobacterial infection in vivo. The method enables direct visualization of early bacterial localization, aggregation, vascular interactions, and macrophage uptake during the initial hours to days following infection, providing a practical foundation for mechanistic studies of host-pathogen interactions under safe experimental conditions.

## Introduction

High-resolution intravital microscopy has transformed the study of dynamic cellular behavior within intact tissues by enabling direct visualization of cell migration, vascular interactions, and immune surveillance at single cell resolution (1–4). While these approaches are widely used in cancer research (3, 5), their application to pulmonary infections has been limited by respiratory motion, tissue fragility and biosafety constraints. Recent advances in thoracic imaging platforms, including vacuum-stabilized windows and permanently implanted optical windows, have addressed many of these technical challenges by providing for stable, motion-free imaging of the pulmonary microenvironment over extended periods (6–10). Permanently implantable windows, in particular, are powerful tools as they enable repeated visualization of vascular flow, immune cell dynamics, and single-cell behavior over hours to weeks, and have been instrumental in defining mechanisms of cancer metastasis, vascular permeability, and immune surveillance (11–13).

Despite these advances, intravital imaging has not previously been applied to *Mycobacterium tuberculosis* (Mtb) infection in the in vivo mammalian lung at single-cell resolution. Prior *in vivo* optical approaches have employed alternative model systems or detection strategies. Real-time imaging of M. marinum infection in optically transparent zebrafish embryos (14) enabled early visualization of mycobacterium-macrophage interactions and granuloma initiation. Intravital two-photon microscopy of BCG-induced granulomas in the mouse liver (15, 16) revealed macrophage and T cell dynamics during granuloma development and demonstrated limited antigen presentation and T cell effector function within granulomatous lesions. Additionally, whole-body optical imaging approaches using fiber-optic microendoscopic excitation (17) and reporter enzyme fluorescence technology (18, 19) have enabled sensitive detection of Mtb in mouse lungs, but cannot detect individual bacilli, track bacterial aggregation dynamics, or visualize direct interactions between mycobacteria and host cells within intact lung tissue. Furthermore, a BSL-3-compatible multiphoton microscopy system has been developed for imaging Mtb in the lung (20), though such setups remain technically demanding and are not widely accessible. Work with virulent Mtb requires BSL-3 containment, posing significant engineering and biosafety challenges for intravital imaging (20). As a result, most studies of early Mtb infection rely on static endpoint assays that cannot capture the spatial and temporal dynamics of bacterial behavior within the lung.

The development of genetically defined, BSL-2-approved triple-auxotrophic Mtb strains provides a unique opportunity to overcome these limitations. The mc^2^7902 strain, a pantothenate-leucine-arginine Mtb auxotroph derivative of H37Rv, is fully attenuated, fails to proliferate in immunocompromised mice, retains classical acid-fast staining, exhibits normal phage susceptibility, and can be safely handled outside of BSL-3 facilities (21). These features make mc^2^7902 an ideal surrogate for establishing the feasibility of intravital imaging approaches and probing early host-pathogen interactions under safer experimental conditions.

Here, we apply the Window for High-Resolution Imaging of the Lung (WHRIL) to visualize fluorescent mc^2^7902::RFP (designated mc^2^8471) in the intact murine lung. Experiments were performed in immunocompromised Rag2^-/-^ mice (22) crossed with MacBlue mice (23), in which the Csf1r promoter drives expression of enhanced cyan fluorescent protein (ECFP) in monocytes, microglia, and subsets of dendritic cells and macrophages. Although the protocol is compatible with a variety of immunocompetent mouse models, with or without fluorescent reporters, the MacBlue transgene enables clear visualization of mononuclear phagocyte populations during early Mtb infection. This platform provides direct, real-time imaging of early Mtb behavior in vivo and establishes a foundation for future mechanistic studies of bacterial physiology, host recognition and immune-mediated clearance.

### Protocols

#### Ethics and Biosafety Statement

All procedures were conducted in accordance with institutional biosafety and animal care guidelines and were approved by the appropriate oversight committees. The strain mc^2^8471 is a pYUB1169 transformant of mc^2^7902, an Mtb strain approved for BSL-2 use. All work involving live bacteria was performed using standard precautions for handling mycobacteria.

##### 1. Construction of the BSL2-approved Td-Tomato-expressing *Mtb* strain mc^2^8471

To enable visualization of Mtb by intravital microscopy, the BSL2-approved triple auxotrophic strain mc^2^7902 was transformed with the episomal mycobacterial plasmid pYUB1169, which contains the Td-Tomato gene expressed from the G13 promoter. Transformation yielded the fluorescent derivative mc^2^8471, which was subsequently used for intravenous infection.

1.1. Generation of Td-Tomato expressing BSL2-safe mc^2^8471 by plasmid transformation
  1.1.1. Inoculate 100 μL mc^2^7902 in 10 mL sterile Middlebrook 7H9 (Difco, Sparks, MD) supplemented with OADC enrichment (10% (v/v), Becton Dickinson), glycerol (0.2 % (v/v)), *D*-pantothenate (24 mg/L), leucine (50 mg/L), arginine (200 mg/L), sodium propionate (0.1 mM), and tyloxapol (0.05% (v/v)) in a 30 mL inkwell bottle.
  1.1.2. Incubate the culture, slowly shaking at 37°C until it has reached an optical density at 600 nm of 0.7-1.0.
  1.1.3. Transfer the cells into a 15 mL conical centrifuge tube and centrifuge 10 min at 3,000 g at room temperature. Discard supernatant and resuspend the cells in 10 mL of sterile 10% glycerol containing 0.05% (v/v) tyloxapol (wash #1). Repeat centrifugation and washes twice. Resuspend the cell pellet in 0.175 ml of 10% glycerol containing 0.05% (v/v) tyloxapol and add 2 µL of pYUB1169 DNA (100-200 ng of DNA).
  1.1.4. Transfer mc^2^7902 and pYUB1169 into a 0.2 cm electroporation cuvette. Wait 10 min and electroporate (BTX Harvard Apparatus, 2500 mV, 1000 ohms, 25 microF).
  1.1.5. Add 1 mL of sterile Middlebrook 7H9 (Difco, Sparks, MD) supplemented with OADC enrichment (10% (v/v)), glycerol (0.2 % (v/v)), *D*-pantothenate (24 mg/L), leucine (50 mg/L), arginine (200 mg/L), sodium propionate (0.1 mM), and tyloxapol (0.05% (v/v)) to the cuvette and transfer the content of the cuvette to a 15 mL conical centrifuge tube. Incubate the tube while shaking at 37°C overnight.
  1.1.6. Plate the transformation on two Middlebrook 7H10 (Difco, Sparks, MD) plates supplemented with OADC enrichment (10% (v/v)), glycerol (0.2 % (v/v)), *D*-pantothenate (24 mg/l), leucine (50 mg/L), arginine (200 mg/L), sodium propionate (0.1 mM), tyloxapol (0.05% (v/v)) and 20 mg/L kanamycin (20 mg/L, Goldbio, St Louis, MO). Incubate the plate at 37°C for three to four weeks. The resulting colonies will be pink.
  1.1.7. Pick one isolated pink colony and grow in 5 mL sterile Middlebrook 7H9 supplemented with OADC enrichment (10% (v/v)), glycerol (0.2 % (v/v)), *D*-pantothenate (24 mg/l), leucine (50 mg/L), arginine (200 mg/L), sodium propionate (0.1 mM), tyloxapol (0.05% (v/v)) and 20 mg/L kanamycin in a 30 mL inkwell bottle.
1.2. Preparation of mc^2^8471 for intravenous infection
  1.2.1. Grow 10 mL of mc^2^8471 in sterile Middlebrook 7H9 supplemented with OADC enrichment (10% (v/v)), glycerol (0.2 % (v/v)), *D*-pantothenate (24 mg/L), leucine (50 mg/L), arginine (200 mg/L), sodium propionate (0.1 mM), tyloxapol (0.05% (v/v)) and 20 mg/l kanamycin in a 30 mL inkwell bottle to mid-log phase (OD_600nm_ should be between 0.7 and 1.0).
  1.2.2. Transfer the cells into a 15 mL conical centrifuge tube and centrifuge 10 min at 3,000 g at room temperature. Discard supernatant and resuspend the cells in 10 mL of sterile Phosphate-buffered saline containing 0.05% (v/v) tyloxapol (PBS-T, wash #1). Repeat centrifugation and washes twice. Resuspend the cell pellet in 4 ml of PBS-T and sonicate the cells (power 100W, 2 x 1000 J).
  1.2.3. Measure the optical density of the culture and dilute it in PBS-T for a final concentration of 5 x 10^6^ CFUs per injection (calculated using the conversion that an OD_600_ of 1 corresponds to 1×10^8^ CFU/ml).

##### 2. Permanent lung window Implantation in mice for intravital imaging

Successful intravital imaging requires surgical implantation of the window for High-Resolution Imaging (WHRIL) to provide stable optical access to the lung while minimizing motion artifacts. This procedure is performed 24 hours before bacterial infection to allow adequate wound healing. The complete detailed protocol for implantation of the WHRIL has been previously published (6). As such, we provide below only a concise summary of the essential steps.

2.1. Induce anesthesia with 5% isoflurane in an induction chamber.
2.2. Administer pre-operative analgesia: inject 10 µL (0.1 mg/kg) of buprenorphine diluted in 90 µL sterile PBS subcutaneously.
2.3. Tie a 2-0 silk suture around the base of a 22G catheter, leaving ~2-inch tails for securing the catheter.
2.4. Intubate the mouse with the suture-tied catheter. Visualize the tracheal opening by trans-illumination and confirm placement using an inflation bulb (bilateral chest rise).
2.5. Connect the catheter to the ventilator.
2.6. Reduce isoflurane to 3% for the remainder of the surgery.
2.7. Remove hair from the right upper thorax using Nair.
2.8. Lift the skin and make a 5-10 mm circular incision ~7 mm left of the sternum and ~7 mm above the subcostal margin (near the intersection of the midline and vertical ear line).
2.9. Grasp and elevate the 6th or 7th rib with forceps and gently pierce the intercostal muscle between them using blunt micro-dissecting scissors (rounded side facing the lung) to enter the thoracic cavity.
2.10. Position the biopsy punch on the cutting tool and guide the cutting tool through the intercostal opening, keeping it parallel to the chest wall. Punch a 5 mm circular defect in the rib cage.
2.11. Place a 5-0 silk purse-string suture ~1 mm from the edge of the hole, interlacing between ribs.
2.12. Align the window frame so the circular chest wall defect fits into the frame groove.
2.13. Gently lift the frame to separate it from the lung and apply a thin layer of cyanoacrylate adhesive to its underside.
2.14. Dry the surface of the lung using a continuous, gentle stream of compressed air. Note: the surface of the lung should appear to gain a matte finish when dry.
2.15. Using positive end expiratory pressure (PEEP), inflate the lung and then adhere the window to the frame. Release the PEEP and keep pressure on the window frame for 20-30 s.
2.16. Place a second 5-0 silk purse-string suture <1 mm from the skin incision edge. Tuck the skin beneath the frame before cinching the purse-string suture tight and then tying securely. Note: ensure that any excess skin is captured under the window frame or trimmed away.
2.17. Using a vacuum pickup, apply a very thin layer of adhesive to the underside of the 5 mm coverslip, scraping off excess on the edge of a rectangular cover glass.
2.18. Again, dry the lung surface with a continuous, gentle stream of compressed air.
2.19. Position the circular coverslip in recess of the window frame, held at an ~15-degree angle from the surface of the lung. Use PEEP to inflate the lung, then establish contact between cover glass and the lung by rotating the coverslip parallel to the lung surface to establish contact. Hold for ~25 s until the adhesive sets.
2.20. Attach a sterile needle to a 1 mL insulin syringe. Insert the needle just below the xiphoid process and advance toward the left shoulder through the diaphragm into the thoracic cavity. Gently aspirate to remove residual intrathoracic air.
2.21. Turn off isoflurane and continue ventilation with 100% oxygen until the mouse begins to recover.
2.22. Administer post-operative analgesia: inject 10 µL (0.1 mg/kg) of buprenorphine diluted in 90 µL sterile PBS subcutaneously.

##### 3. Intravital imaging

Following window implantation, mice require a recovery period before imaging. Optimal imaging quality is achieved 1 to 5 days post-surgery, allowing visualization of early Mtb infection dynamics through repeated imaging sessions.

3.1. Induce anesthesia with 3% isoflurane, then maintain at 1.5% for the duration of imaging.
3.2. Maintain body temperature at ~37 °C using a forced air heated chamber with temperature sensor feedback loop.
3.3. Insert the custom stage plate-window frame adapter between the skin and the outer rim of the lung window frame and secure to a standard microscope stage plate using adhesive tape.
3.4. Place the stage plate on the microscope translation stage.
3.5. Image macrophages (CFP), vasculature (FITC-dextran), and Mtb (tdTomato) using an Olympus FluoView 1000 on an inverted IX81 microscope with a 25× 1.05 NA objective.
3.6. Inject FITC-dextran (150 kDa; 20 mg/mL, 100 µL) retro-orbitally while the mouse is on the stage.
3.7. Acquire snapshot, z-stack, or time-lapse images of the lung vasculature before introduction of tuberculosis using the following parameters:
  a. Image depth: 12-bit
  b. Frame rate: 2 fps
  c. Excitation: 405 nm, 488 nm, 559 nm
  d. Detection filters: 430–470 nm, 505–540 nm, 575–675 nm
3.8. Retro-orbitally inject 5 x 10^6^ CFUs tdTomato-expressing mc^2^8471.
3.9. Acquire snapshot, z-stack, and time-lapse images of tuberculosis and fluorescently labeled cells and vasculature.
3.10. After imaging, turn off isoflurane and remove the mouse from the microscope. Keep the mouse warm and monitor until full recovery.
3.11. Repeat imaging at specific timepoints. For each imaging session, re-inject FITC-dextran (100 µL, retro-orbitally) to visualize vasculature, then acquire images as described in steps 3.8-3.10.

### Representative Results

Following electroporation and kanamycin selection, isolated colonies of mc^2^8471 exhibited bright red fluorescence, confirming stable tdTomato expression. Fluorescent bacteria were expanded, prepared as a single-cell suspension and administered intravenously to mice bearing surgically implanted WHRIL. Intravital imaging was performed to visualize early Mtb infection dynamics in the lung microenvironment. For this study, a series of intravital images were acquired within 60 minutes, at 6 hours, and 3 days after mc^2^8471 injection into a Rag2^-/-^ x Macblue mouse (**Figures 1–3)**. Within a few minutes following mc^2^8471 intravenous infection, bacteria could be observed within blood vessels, illustrating how rapidly Mtb reaches and localizes within the lung vasculature (**Figure 1**). One hour post Mtb infection, the bacilli can be seen still located near the blood vessels. They appear either as single cells or as an organized cluster of bacteria (**Figure 2A**).

**Figure 1:**
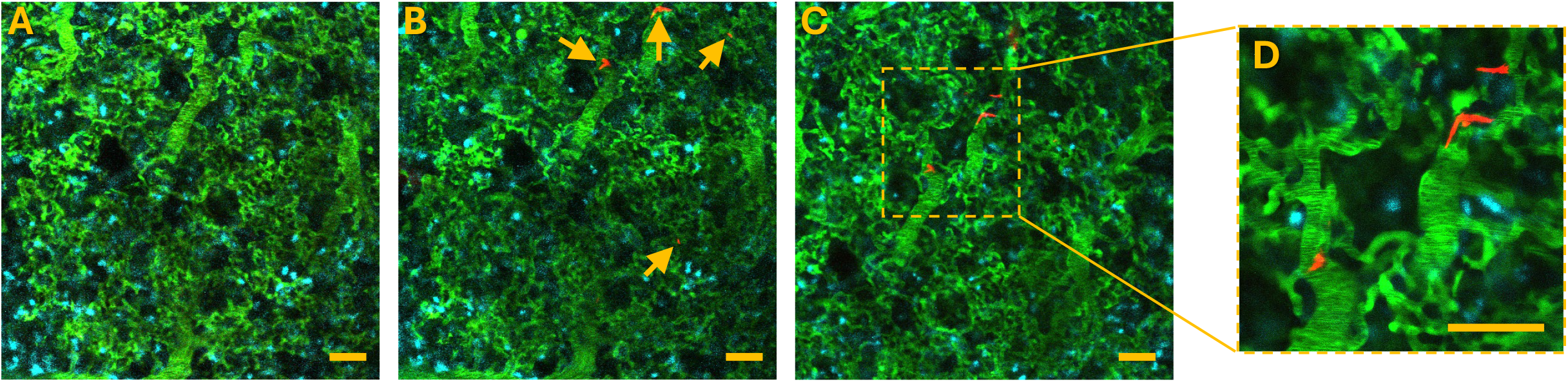
In vivo imaging of lung vasculature before and immediately after injection of mc^2^8471. **(A, B)** In vivo images of blood vessels (green) before (A) and immediately after (B) intravenous injection of mc^2^8471. White arrows in (B) indicate the appearance of mc^2^8471 (red) following injection. **(C)** Representative image showing mc^2^8471 within lung blood vessels. **(D)** Magnified view of the inset region from (C). Green = FITC-labeled 150 kDa Dextran. Red = tdTomato mc^2^8471 Mtb. Cyan = CFP-labeled macrophages. Scale bars = 50 μm.

**Figure 2:**
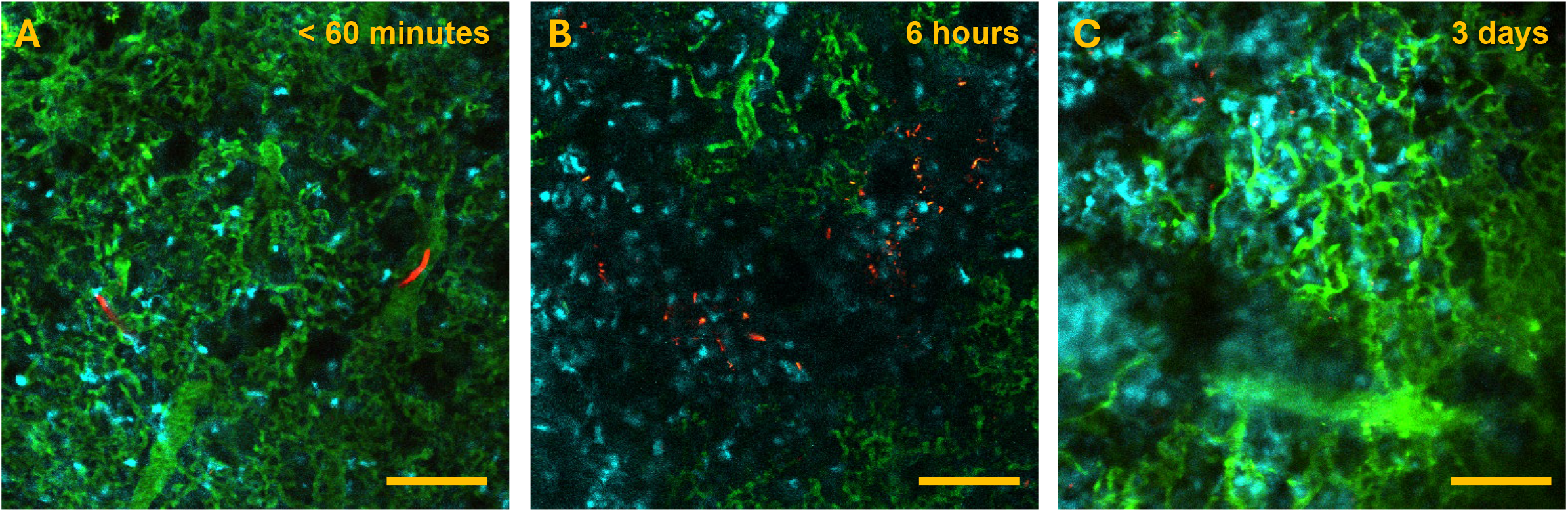
Dynamics of mc^2^8471 localization from time of injection to 3 days post-injection. **(A)** Early localization of mc^2^8471 within 60 minutes after intravenous injection. **(B)** In another field of view at 6 hours post-injection, mc^2^8471 forms aggregates, leading to blood vessel occlusion. **(C)** By 3 days post-injection, most Mtb bacilli have cleared; however, some individual Mtb bacilli remain and are taken up by macrophages. Green = FITC-labeled 150 kDa Dextran. Red = tdTomato mc^2^8471 Mtb. Cyan = CFP-labeled macrophages. Scale bars = 100 μm.

By six hours post-Mtb infection, a drastic change in bacteria localization was observed, with bacilli found widely disseminated throughout the lung parenchyma in regions of the lungs where no macrophages or blood vessels could be visualized (**Figure 2B**), indicating complete vascular occlusion and loss of local perfusion.

By three days post-infection (**Figure 3**), Mtb bacilli were markedly reduced compared to the six-hour timepoint, indicating substantial bacterial clearance from the lung during this period. When detected, bacteria were found predominantly within macrophages, with multiple bacilli per cell (multiplicity of infection greater than one). This shift from extracellular aggregates at six hours to predominantly intracellular bacteria by day three demonstrates successful phagocytic uptake and suggests that the bacilli replication cannot outpace immune clearance at this early stage.

**Figure 3:**
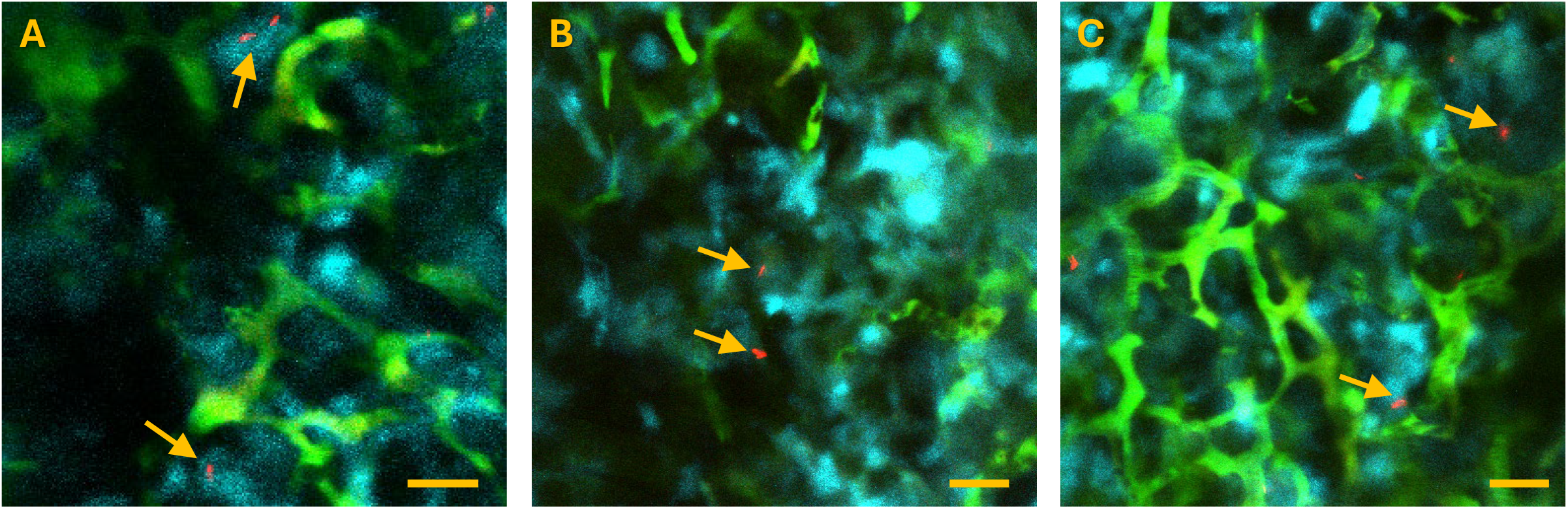
Uptake of mc^2^8471 by macrophages at 3 days post-injection. **(A–C)** Representative images showing Mtb internalized by macrophages. Green = FITC-labeled 150 kDa Dextran. Red s= tdTomato mc^2^8471 Mtb. Cyan = CFP-labeled macrophages. Scale bars = 20 μm.

Together, these observations demonstrate that the WHRIL platform enables direct visualization of sequential early events in Mtb infection, including vascular delivery, aggregation, parenchymal dissemination, and macrophage uptake, at single-cell resolution in vivo.

## Discussion

This work establishes two key advances. First, it demonstrates that intravital lung imaging can be successfully adapted to visualize Mtb behavior at single-cell resolution in intact lung tissue building upon pioneering intravital imaging studies of mycobacterial infections in zebrafish (14) and BCG granulomas in mouse liver(15, 16), while extending these approaches to enable direct visualization of individual bacilli and their interactions with host cells in the mammalian lung. Second, by leveraging a genetically defined, BSL-2–approved attenuated Mtb strain, this approach overcomes longstanding biosafety and technical barriers that have prevented real-time visualization of mycobacterial infection in vivo, making intravital imaging of Mtb broadly accessible.

By integrating the Window for High-Resolution Imaging of the Lung (WHRIL) with the fluorescent derivative of the triple-auxotrophic strain mc^2^7902, we directly observed early infection events that were previously inferred only indirectly. These include rapid vascular delivery of bacilli to the lung, early aggregation and vascular occlusion, dissemination into the lung parenchyma, and subsequent uptake by pulmonary macrophages. Importantly, because these events were captured dynamically within the same tissue environment, this method reveals spatial and temporal relationships not accessible through conventional endpoint assays.

Although this study employs intravenous inoculation of an attenuated strain rather than aerosol delivery of virulent Mtb, the biological behaviors observed (vascular arrest, aggregation, parenchymal dissemination, and phagocyte uptake) mirror key early steps thought to occur during pulmonary infection. Intravenous inoculation provides a controlled and reproducible system to study early lung host–pathogen interactions under BSL-2 conditions, enabling mechanistic insights that would otherwise require highly specialized BSL-3 imaging infrastructure. Extending intravital lung imaging to aerosol infection with virulent Mtb under BSL-3 conditions requires specialized containment infrastructure(20) and remains an important future direction.

Our approach is distinct from prior *in vivo* imaging strategies for tuberculosis. Whole-body optical detection methods using fiber-optic microendoscopic excitation(17) achieved ~100 CFU detection sensitivity in the lung by delivering intravital illumination via tracheal catheter to excite BlaC-activated near-infrared fluorescent substrates. Similarly, reporter enzyme fluorescence imaging(18) and near-infrared fluorescent protein-expressing Mtb(19) enabled longitudinal monitoring of bacterial burden and drug responses in live animals. While these technologies provide sensitive quantification of pulmonary infection, they measure aggregate bacterial loads and do not visualize individual bacilli, distinguish single bacteria from clusters, or reveal spatial relationships between mycobacteria and host cells at cellular resolution within intact tissue architecture. Furthermore, although BSL-3-compatible multiphoton systems have demonstrated proof-of-principle for imaging virulent Mtb in the lung(20), such implementations require specialized containment infrastructure and remain inaccessible to most laboratories. By combining a permanent lung window with BSL-2-approved, fluorescent triple-auxotrophic Mtb, our platform achieves detection and tracking of individual fluorescent bacilli over multiple days within the intact lung microenvironment under standard BSL-2 conditions.

Beyond establishing technical feasibility, this BSL-2 intravital imaging platform enables experimental investigations that were previously inaccessible. Mtb undergoes profound metabolic remodeling during infection, including shifts toward host lipid and cholesterol utilization (24), yet how these physiological states influence bacterial morphology, aggregation, trafficking, or immune recognition in vivo remains unclear. Because this system permits real-time visualization of individual bacilli within intact lung tissue, it can be directly applied to assess how defined metabolic conditions or growth states alter early host–pathogen interactions in the lung.

Similarly, this platform enables systematic analysis of genetically defined mutants with altered cell-envelope or virulence properties. These include acid-fast–deficient mutants (25), non-cording variants that may influence aggregation or vascular occlusion, ESX-1–deficient strains resembling BCG (26), and other mutants affecting persistence or immune evasion. Since mc^2^7902-derived strains remain BSL-2 compatible (21), they can be imaged repeatedly at high resolution, allowing direct dissection of how specific bacterial attributes shape early infection dynamics in vivo.

A particularly compelling future application of this platform is the direct visualization of immune-mediated killing of Mtb. The conditionally persistent mutant mc^2^7901, which persists in immunodeficient hosts but is sterilized by adaptive immunity in immunocompetent mice (27), provides a uniquely tractable substrate for such studies. Combining aerosol infection of fluorescent mc^2^7901 with intravital lung imaging could enable the real-time visualization of when and where immune-mediated sterilization occurs at the single-cell level.

Critically, this system also enables direct comparison of early bacillary fate in unimmunized mice versus mice immunized with mc^2^7902, a regimen known to induce exceptionally strong protection (28). Such studies could reveal whether sterilization correlates with altered macrophage behavior, accelerated intracellular killing, or changes in spatial organization within the lung, thereby distinguishing immune mechanisms that restrain bacterial growth from those that achieve true clearance.

In summary, this work provides the first demonstration that intravital lung imaging can be successfully adapted to study mycobacterial infection in vivo under BSL-2 conditions, providing a practical platform to probe early bacterial physiology, host recognition, and immune responses in real time, while laying essential groundwork for future studies with virulent Mtb. By enabling direct visualization of early infection dynamics—and prospectively, immune-mediated sterilization—this approach opens experimental avenues that have previously been inaccessible in tuberculosis research.

### Troubleshooting and Limitations

Successful application of this intravital imaging protocol depends on both surgical technique and careful preparation of fluorescent Mtb. Motion artifacts during imaging are the result of suboptimal lung window implantation, reflecting incomplete stabilization of the lung surface. Inadequate sealing of the coverslip will be immediately evident as mice will be unable to recover from the surgery. Ensuring complete removal of intrathoracic air, proper use of positive end-expiratory pressure during window attachment, and gentle drying of the lung surface before adhesion are critical steps for achieving stable, high-resolution imaging.

Preparation of a single-cell bacterial suspension is also essential. Incomplete dispersion of mc^2^8471 can lead to artifactual aggregation prior to injection, complicating interpretation of early in vivo clustering events. Adequate sonication and inclusion of tyloxapol during washing steps minimize clumping and improve reproducibility. Fluorescence intensity should be verified prior to infection, as reduced td-Tomato signal may indicate plasmid loss or suboptimal growth conditions.

A key limitation of the current protocol is the use of an attenuated, triple-auxotrophic Mtb strain delivered intravenously. Although this approach enables imaging under BSL-2 conditions and reveals biologically informative early events, including vascular arrest, aggregation, dissemination, and macrophage uptake, it does not fully recapitulate all aspects of aerosol infection with virulent Mtb. As such, quantitative features such as dissemination kinetics and immune cell recruitment should be interpreted within the constraints of this model. Extending intravital lung imaging to aerosol infection with virulent Mtb under BSL-3 conditions remains an important long-term goal that will require substantial engineering and containment innovations, though proof-of-principle BSL-3 multiphoton systems(20) suggest such adaptations are feasible.

Finally, the WHRIL platform provides optical access to a defined region of the lung and therefore captures a spatially restricted view of infection. While repeated imaging allows longitudinal analysis within the same microenvironment, care should be taken when generalizing observations across the entire lung. Despite these limitations, this protocol offers a unique and powerful approach to directly visualize early mycobacterial behavior and host-pathogen interactions in vivo, providing insights not achievable through conventional endpoint assays.

## Acknowledgements

This work was supported by the following grants: AI26170; The Evelyn-Lipper Family Foundation and the Gruss Lipper Biophotonics Center.

## Table of Materials

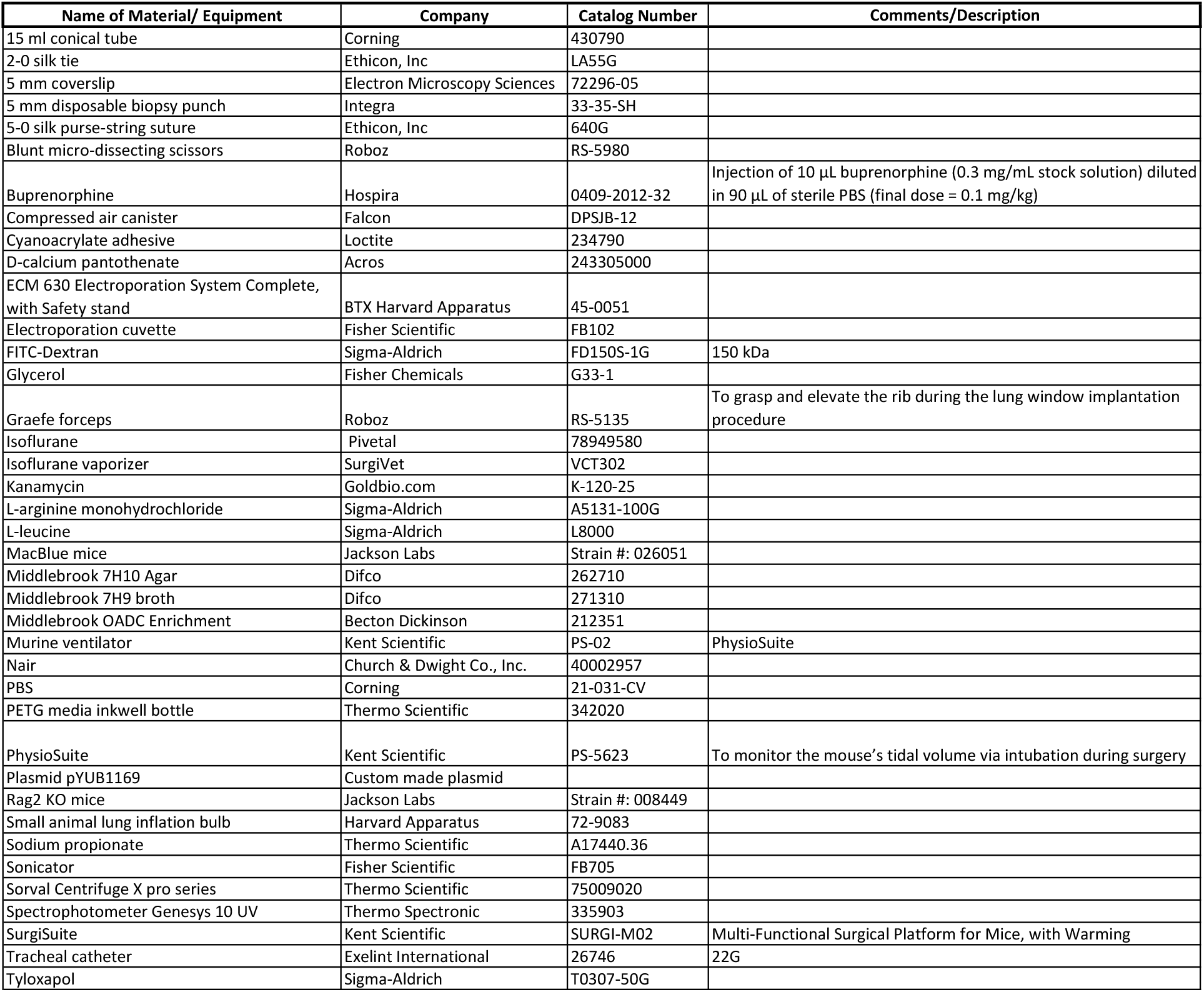
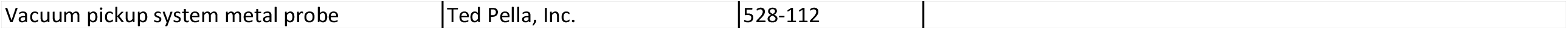

## Notes

### Competing Interest Statement

The authors have declared no competing interest.

## References

1. Scheele C, Herrmann D, Yamashita E, Celso CL, Jenne CN, Oktay MH, et al. Multiphoton intravital microscopy of rodents. Nat Rev Methods Primers. 2022;2(1):1–26.

2. Niesner RA, Hauser AE, Entenberg D. Life Through a Lens: Technological Development and Applications in Intravital Microscopy. Cytometry A. 2020;97(5):445–7.

3. Perrin L, Bayarmagnai B, Gligorijevic B. Frontiers in Intravital Multiphoton Microscopy of Cancer. Cancer Rep (Hoboken). 2020;3(1):e1192.

4. Weigert R, Porat-Shliom N, Amornphimoltham P. Imaging cell biology in live animals: ready for prime time. J Cell Biol. 2013;201(7):969–79.

5. Entenberg D, Oktay MH, Condeelis JS. Intravital imaging to study cancer progression and metastasis. Nat Rev Cancer. 2023;23(1):25–42.

6. Borriello L, Traub B, Coste A, Oktay MH, Entenberg D. A Permanent Window for Investigating Cancer Metastasis to the Lung. J Vis Exp. 2021(173):e62761.

7. Rodriguez-Tirado C, Kitamura T, Kato Y, Pollard JW, Condeelis JS, Entenberg D. Long-term High-Resolution Intravital Microscopy in the Lung with a Vacuum Stabilized Imaging Window. J Vis Exp. 2016(116):e54603.

8. Presson RG, Jr., Brown MB, Fisher AJ, Sandoval RM, Dunn KW, Lorenz KS, et al. Two-photon imaging within the murine thorax without respiratory and cardiac motion artifact. Am J Pathol. 2011;179(1):75–82.

9. Looney MR, Thornton EE, Sen D, Lamm WJ, Glenny RW, Krummel MF. Stabilized imaging of immune surveillance in the mouse lung. Nat Methods. 2011;8(1):91–6.

10. Entenberg D, Voiculescu S, Guo P, Borriello L, Wang Y, Karagiannis GS, et al. A permanent window for the murine lung enables high-resolution imaging of cancer metastasis. Nat Methods. 2018;15(1):73–80.

11. Alieva M, Ritsma L, Giedt RJ, Weissleder R, van Rheenen J. Imaging windows for long-term intravital imaging: General overview and technical insights. Intravital. 2014;3(2):e29917.

12. Prunier C, Chen N, Ritsma L, Vrisekoop N. Procedures and applications of long-term intravital microscopy. Methods. 2017;128:52–64.

13. Coste A, Oktay MH, Condeelis JS, Entenberg D. Intravital Imaging Techniques for Biomedical and Clinical Research. Cytometry A. 2020;97(5):448–57.

14. Davis JM, Clay H, Lewis JL, Ghori N, Herbomel P, Ramakrishnan L. Real-time visualization of mycobacterium-macrophage interactions leading to initiation of granuloma formation in zebrafish embryos. Immunity. 2002;17(6):693–702.

15. Egen JG, Rothfuchs AG, Feng CG, Horwitz MA, Sher A, Germain RN. Intravital imaging reveals limited antigen presentation and T cell effector function in mycobacterial granulomas. Immunity. 2011;34(5):807–19.

16. Egen JG, Rothfuchs AG, Feng CG, Winter N, Sher A, Germain RN. Macrophage and T cell dynamics during the development and disintegration of mycobacterial granulomas. Immunity. 2008;28(2):271–84.

17. Nooshabadi F, Yang HJ, Cheng Y, Durkee MS, Xie H, Rao J, et al. Intravital excitation increases detection sensitivity for pulmonary tuberculosis by whole-body imaging with beta-lactamase reporter enzyme fluorescence. J Biophotonics. 2017;10(6-7):821–9.

18. Sharan R, Yang HJ, Sule P, Cirillo JD. Imaging Mycobacterium tuberculosis in Mice with Reporter Enzyme Fluorescence. J Vis Exp. 2018(132):e56801.

19. Sommer R, Cole ST. Monitoring Tuberculosis Drug Activity in Live Animals by Near-Infrared Fluorescence Imaging. Antimicrob Agents Chemother. 2019;63(12):e01280–19.

20. Barlerin D, Bessiere G, Domingues J, Schuette M, Feuillet C, Peixoto A. Biosafety Level 3 setup for multiphoton microscopy in vivo. Sci Rep. 2017;7(1):571.

21. Vilcheze C, Copeland J, Keiser TL, Weisbrod T, Washington J, Jain P, et al. Rational Design of Biosafety Level 2-Approved, Multidrug-Resistant Strains of Mycobacterium tuberculosis through Nutrient Auxotrophy. mBio. 2018;9(3):e00938–18.

22. Shinkai Y, Rathbun G, Lam KP, Oltz EM, Stewart V, Mendelsohn M, et al. RAG-2-deficient mice lack mature lymphocytes owing to inability to initiate V(D)J rearrangement. Cell. 1992;68(5):855–67.

23. Ovchinnikov DA, van Zuylen WJ, DeBats CE, Alexander KA, Kellie S, Hume DA. Expression of Gal4-dependent transgenes in cells of the mononuclear phagocyte system labeled with enhanced cyan fluorescent protein using Csf1r-Gal4VP16/UAS-ECFP double-transgenic mice. J Leukoc Biol. 2008;83(2):430–3.

24. Pandey AK, Sassetti CM. Mycobacterial persistence requires the utilization of host cholesterol. Proc Natl Acad Sci U S A. 2008;105(11):4376–80.

25. Bhatt A, Fujiwara N, Bhatt K, Gurcha SS, Kremer L, Chen B, et al. Deletion of kasB in Mycobacterium tuberculosis causes loss of acid-fastness and subclinical latent tuberculosis in immunocompetent mice. Proc Natl Acad Sci U S A. 2007;104(12):5157–62.

26. Hsu T, Hingley-Wilson SM, Chen B, Chen M, Dai AZ, Morin PM, et al. The primary mechanism of attenuation of bacillus Calmette-Guerin is a loss of secreted lytic function required for invasion of lung interstitial tissue. Proc Natl Acad Sci U S A. 2003;100(21):12420–5.

27. Vilcheze C, Porcelli SA, Chan J, Jacobs WR, Jr. Sterilization by Adaptive Immunity of a Conditionally Persistent Mutant of Mycobacterium tuberculosis. mBio. 2021;12(1);e02391–20.

28. Vilcheze C, Rajagopalan S, Jacobs Jr WR. Intravenous Immunization with Triple Auxotrophs of Mycobacterium tuberculosis: A novel vaccine strategy against tuberculosis. bioRxiv. 2024:2024.05.15.594337.

